# Combinatorial effects of bovicin HC5 and epsilon-polylysine against *Alicyclobacillus acidoterrestris* in orange juice

**DOI:** 10.1101/2025.01.11.632513

**Authors:** Nneka V Iduu

## Abstract

**BACKGROUND:** *Alicyclobacillus acidoterrestris*, a spoilage-causing bacterium in fruit juices remains a concern post-pasteurization, making antimicrobial peptides (AMPs) a promising alternative for its control. Bovicin HC5 and epsilon-polylysine were tested for their antimicrobial efficacy against *A. acidoterrestris* DSMZ 2498 in both AAM broth and orange juice. This study determined the minimum inhibitory concentrations (MICs) and minimum bactericidal concentration (MBC) for epsilon-polylysine and bovicin HC5 against 10^6^ CFU/mL vegetative cells and activated endospores of *A. acidoterrestris* DSMZ 2498. Their combined effect was also assessed using the checkerboard method. Viable cell counts were measured after 48 hours at 45 °C, and thermal resistance of endospores was examined after exposure to both AMPs as well as their combinations, by determining the exposure time (in minutes) required to kill 90 % of the initial population at 95 °C. Finally, atomic force microscopy (AFM) was employed to investigate their impact on the cell structure and morphology of *A. acidoterrestris*.

**RESULTS:** Bovicin HC5 and epsilon-polylysine demonstrated antibacterial effects against *A. acidoterrestris* DSMZ 2498 inoculated in orange juice. The combination of both AMPs exhibited an additive effect, significantly reducing the number of viable cells after a 48-hour incubation at 45 °C and a 90 % reduction in thermal resistance at 95 °C. Atomic force microscopy (AFM) revealed structural and morphological changes in *A. acidoterrestris* cells treated by these AMPs.

**CONCLUSIONS:** The findings in this study revealed the potential application of bovicin HC5 and epsilon-polylysineas natural preservatives in hurdle technologies to control *A. acidoterrestris* and improve the microbiological stability and safety of fruit juices.

## INTRODUCTION

Commercial fruits are subjected to pasteurization because they are suitable substrates for the growth of spoilage microorganisms^1^. Despite pasteurization, the survival of spores, especially those of thermoacidophilic bacteria, and their subsequent germination and growth into vegetative cells can occur in low-pH juices, contributing to food product spoilage^2^. *A. acidoterrestris*, a Gram-positive, non-pathogenic, thermoacidophilic, strictly aerobic bacterium is considered an important spore-forming microorganism that causes the deterioration of juices^3,4^. *A. acidoterrestris* produces substances that promote odors in juices, such as guaiacol, 2,6- dibromophenol, 2,6-dichlorophenol, and halophenols, compromising the quality of the products^5^. It also grows at temperatures ranging from 26 to 60 °C, and a pH ranging from 2.0 to 6.0 allowing its survival during commercial pasteurization process which makes this bacterium a key quality control target for pasteurized fruit juices and relevant beverages^4,6^ .

Various physical treatments like high hydrostatic pressure, ultra-high-pressure homogenization, UV-C light inactivation, irradiation, microwaves, ultrasonic waves, and ohmic heating have been explored to control *A. acidoterrestris*^7,8^. While these methods have shown effectiveness, some drawbacks include limited penetration into food matrices, absence of sporicidal activity, off-odors, low consumer acceptance, and adverse impacts on organoleptic properties^7^.

In recent years, there has been an increasing interest in the use natural additives such as antimicrobial peptides to preserve food products due to their ability to retain food quality without harmful effects^9^.

Epsilon-polylysine is a natural antimicrobial cationic peptide produced by *Streptomyces albulus* ssp. *lysinopolymerus* that is generally regarded as safe (GRAS) as a food preservative^10,11^. Epsilon-Polylysine is characterized as being edible, non-toxic to humans, water-soluble, and stable at high temperatures ^10,12^. This compound exhibits a wide antimicrobial spectrum against bacteria, yeasts, and molds, and has been reported to have antibacterial effects on *A. acidoterrestris* ^10,6,13^.

Bacteriocins are ribosomally synthesized and extracellularly-released antimicrobial peptides produced by bacteria that vary in structure, biochemical properties, mode of action, and spectrum of activity^14^. Bovicin HC5, a bacteriocin from *Streptococcus equinus HC5*, is very stable at high temperatures and in acidic environments making it suitable to control thermoacidophilic spoilage bacteria^15^. It inhibits several Gram-positive bacteria, exhibiting bactericidal and sporicidal activity against several strains of food-borne pathogenic and spoilage microorganisms such as *A. acidoterrestris* in fruit juices and pulps, hence, is suggested as an additive for food preservation ^14,16,1^.

Since the individual antibacterial activity of bovicin HC5 and epsilon-polylysine against *A. acidoterretris* has been demonstrated in previous works, we hypothesized that their use in combination could improve their effect against *A. acidoterrestris* and the efficacy of thermal processing of fruit juices. Thus, this work evaluated the combined effect of bovicin HC5 and epsilon-polylysine against vegetative cells and germinated endospores of *A. acidoterrestris* in orange juice focusing on improving the prevention of fruit juice spoilage.

## MATERIALS AND METHODS

### Media preparation, bacteria strains, and culture conditions

A semi-synthetic basal medium (PC) prepared under the flux of oxygen (O_2_)-free carbon dioxide flow was used to cultivate *Streptococcus equinus* HC5 anaerobically at 39 °C. The medium contained: 0.292 g/L KH_2_PO_4_, 0.480 g/L (NH4)_2_SO_4_, 0.480 g/L NaCl, 0.100 g/L MgSO_4_·7H_2_O, 0.640 g/L, CaCl_2_·2H_2_O, 0.500 g/L cysteine hydrochloride, 1 g/L Trypticase (BBL Microbiology Systems, Cockeysville, MD, USA), 0.500 g/L yeast extract, 4g of Na_2_CO_3_, and 8 g/L glucose.

The medium pH was adjusted to 6.5 with sodium hydroxide (NaOH, 3 mol/L) before sterilization (121 °C, 15 min). Bacterial growth (OD600nm) was monitored using a Spectronic 200E spectrophotometer (Thermo Fisher Scientific, Madison, WI, USA).

*A. acidocaldarius* medium (AAM) was used to culture and enumerate *A. acidoterrestris* DSMZ 2498 under aerobic conditions at 45 °C. The medium was composed of solution A (0.25 g CaCl_2_·7H_2_O, 0.5 g MgSO_4_·7H_2_O, 0.2 g (NH4)_2_SO_4_, 3.0 g KH_2_PO_4_); yeast extract (2.0 g); glucose (5.0 g). The volume of the solution was adjusted to 1000 mL (for liquid medium) and 500 mL (for solid medium) with distilled water and the pH was adjusted to 4.0 with 1 mol/L hydrochloric acid (HCL). Solution B contained 0.1 g ZnSO_4_·7H_2_O; 0.03 g MnCl_2_·4H_2_O; 0.3 g H_3_BO_3_; 0.2 g CoCl_2_·6H_2_O; 0.01 g CuCl_2_·2H_2_O; 0.02 g NiCl_2_·6H_2_O, and 0.03 g NaMoO_4_·2H_2_O, with volume adjusted to 1000 mL distilled water. Solution C was composed of 15 g agar prepared with 500 mL distilled water. All solutions were sterilized separately (121°C⁄ 15 min). Solution B was sterilized through a 0.22 μm membrane (Millipore 67120; Molsheim, France). After sterilization, 1 mL of the trace-element solution SL-6 (solution B) was added to solution A (with 1000 mL distilled water) for liquid medium, and also added to solution A (with 500 mL distilled water) combined with solution C for solid medium.

### Preparation of *A. acidoterrestris* DSMZ 2498 spore suspensions

Endospores of *A. acidoterrestris* DSMZ 2498 were obtained in AAM broth as described by Silva et al.^17^ with modifications^1^. The culture was incubated at 45 °C for 120 h, under orbital agitation (120 rpm), and subsequently refrigerated at 4 °C for 48 h. Slides were prepared for microscopic observation of spores after malachite green dye staining. The culture was then centrifuged (9000 × g, 10 min, 5 °C) (4 K15, D-37520; Sigma 4K15, SIGMA Laborzentrifugen GmbH, Osterode, Germany). The supernatant was discarded, and the pellet was suspended in 20 mL of phosphate buffer (pH 4.0, 50 mmol/L). The suspension was then incubated in a thermostatically controlled water bath (Thermomix. BM-S, B. Braun Biotech International, Melsungen, Germany) at 80 °C for 10 min to activate the endospores. After heat treatment, the endospore suspension was centrifuged three times (9000 × g, 10 min, 5 °C), and the supernatant was discarded. The pellet was then re-suspended in 10 mL of phosphate buffer (pH 4.0, 50 mmol/L) and stored at 4 °C until use. The endospores were enumerated by counting the colonies resulting from viable spores that germinated in AAM media.

### Preparation of antimicrobial peptides (AMPs)

A commercial powdered epsilon-polylysine from Zhengzhou Bainafo Bioengineering Co., Ltd (Zhengzhou, China) was used in this study. The stock solution was prepared in sterile distilled water, then filter sterilized (pore size, 0.22 μm), and prepared fresh before use. A series of 2-fold dilutions was also prepared using sterilized distilled water^6^.

Extracts of bovicin HC5 were prepared as described by Mantovani et al.^18^. The pH of the culture was adjusted to 7 followed by a heat treatment at 75 °C for 30 minutes. Stationary-phase *S. equinus* HC5 was harvested by centrifugation (10,000 x g) for 20 min and the supernatant was discarded. The weight of the wet cell mass was recorded. The cell pellet was suspended in 0.1 M acidic NaCl solution (50 mL, pH 2.0) and left overnight at 4 °C with continuous stirring. The suspensions were then centrifuged (10,000 x g) to remove cells and recover the crude extract for purification. Purification of bovicin HC5 was performed using a Sep-Pak C18 Plus Short Cartridge column as the stationary phase; a 0.1 % formic acid solution was used as the mobile phase and elution was carried out with 50 % acetonitrile. Then the purified cell-free supernatant was lyophilized (Edwards; Super Modulyo, Livermore, CA, USA). Serial dilutions were made using sterilized distilled water. The titer of epsilon-polylysine and bovicin HC5 were expressed in terms of microgram per milliliter (μg/mL).

### Determination of the AMPs activity

The antimicrobial activity was analyzed by agar well diffusion assay using *A. acidoterrestris* DSMZ 2498 as the target bacteria. Aliquots of 25 μL of each dilution of the AMPs were applied into wells that were perforated aseptically using sterile plastic straws with a diameter of 5 mm in AAM medium previously inoculated with *A. acidoterrestris* DSMZ 2498 (10^6^ CFU/mL). The agar plates were then incubated at 4 °C overnight to allow the peptides to diffuse and then the plates were incubated at 45 °C for 24 h to allow the growth of the indicator organism.

Antimicrobial activity was detected by measuring the zone of inhibition (including the diameter of the wells) that appeared after the incubation period.

### Determination of the Minimum inhibitory and bactericidal concentrations (MICs and MBCs) of the AMPs

The antimicrobial efficacies of epsilon-polylysine and bovicin HC5 were tested by measuring their MICs against *A. acidoterrestris* DSMZ 2498 in AAM broth. The MICs were determined in 96-well microdilution plates, according to M7-A6 204 ^19^. A series of two-fold dilutions of bovicin HC5 was prepared using sterilized distilled water, then 20 μL of these dilutions were dispensed in wells of a 96-well plate containing a mixture of 180 μL of AAM broth (pH 4) and approximately 10^6^ CFU/mL of vegetative cells of *A. acidoterrestris* DSMZ 2498. Similarly, epsilon-polylysine was diluted and tested as described for bovicin HC5 treatment.

Concentrations of bovicin HC5 and epsilon-polylysine suitable for testing their MIC were determined based on preliminary results obtained from its antimicrobial activity. Final concentrations of bovicin HC5 used were 0.31, 0.63, 1.25, 2.50, 5.00 and 10.00 μg/mL. For epsilon-polylysine, the following concentrations were used 3.91, 7.81, 15.63, 31.25, 62.50, and 125.00 μg/mL. The 96-well plates were incubated at 45 °C for 24 h and bacterial growth was measured at 600 nm in a Spectronic 20D+ (Thermal Electron, Madison, WI, USA). The specific growth rate, lag phase, and OD values after 24 h incubation was determined^15^.

The same protocol was repeated to determine the minimum inhibitory concentrations (MICs) against *A. acidoterrestris* DSMZ 2498 spore cells. Prior to incubation at 45 °C for 24 h, the 96 well plates were heat shocked at 80 °C for 10 min to activate dormant endospores After this period, the Minimum Bactericidal Concentration (MBC) was determined by sub-culturing 20 μL from each well with no growth on AAM agar and incubating at 45 °C for a further 24 h. The absence of growth was the parameter used to define if a given concentration was bactericidal against *A. acidoterrestris*. The tests were performed in technical triplicate and two biological replicates.

### Analysis of the combined effect of the AMPs against *A. acidoterrestris* DSMZ 2498 in AAM broth

The checkerboard method was used to evaluate the antibacterial effect of the combination of bovicin HC5 and epsilon-polylysine using 96-well plates to calculate the Fractional Inhibitory Concentration (FIC) index^20^. The test was performed using different combined concentrations, based on the MIC results of each antimicrobial peptide^4^. The concentrations of each AMP used were 4X, 2X, 1X, 1/2X, 1/4X, and 1/8X of their MIC. Aliquots of 20 uL of each AMP were added to 160 μL of AAM broth containing approximately 10^6^ CFU/mL of *A. acidoterrestris* (vegetative cells) in the wells of a 96-well plate and the microplate was incubated at 45 °C for 24 h. Similarly, bacterial growth was monitored via changes in optical density (OD) at 600 nm in a Spectronic 20D+ (Thermal Electron, Madison, WI, USA)^15^. The same protocol was repeated using endospores of *A. acidoterrestris* DSMZ as inoculum. Prior to incubation at 45 °C for 24 h, the 96-well plates were heat shocked at 80 °C for 10 min to activate dormant endospores.

The FIC indexes were calculated as FIC_A_+FIC_B_, where FIC_A_ = MIC_A_ combined/MIC_A_ alone and FIC_B_ = MIC_B_ combined/MIC_B_ alone. Results were interpreted as synergism (FIC ≤ 0.5), addition (0.5 < FIC ≤ 1), indifference (1 < FIC ≤ 4) or antagonism (FIC > 4)^20^. The assay was performed in technical and biological duplicates.

### Time-kill assay of the AMPs alone and in combination in orange juice

Commercial shelf-stable orange juice (pH 3.68, TSS 11 °Bx, sugar content 17 g, carbohydrate 21 g, protein 1.4 g, vitamin C 45 mg) was obtained from a local grocery store in Vicosa, MG, Brazil. The juice was analyzed for the absence of vegetative cells or spores according to Anjos et al.^21^. The soluble solid content and pH of the juice were determined. Aliquots of the orange juice were transferred into glass tubes, autoclaved, and cooled to room temperature. The concentration of AMPs used was determined by preliminary testing. The assay was performed by adding 0.5 mL of the serially diluted AMPs into 4.5 mL of orange juice containing approximately 10^6^ CFU/mL of *A. acidoterrestris* (vegetative cells) to give the following final concentrations of bovicin HC5: 5, 10, 20, 40, 80 μg/mL and 7.81, 15.63, 31.25, 62.5, 125.0, and 250.0 μg/mL of epsilon-polylysine. When the AMPs were tested in combination, 4 mL of orange juice was used to achieve combined ratios of bovicin HC5:epsilon-polylysine; 5.0:7.81, 10.0:15.63, 20.0:31.25, 40.0:62.5, 80.0:125.0 μg/mL. All tubes were incubated at 45 °C in a water bath for 0, 12, 24, 36, and 48 h. Then 100 μL was withdrawn from each tube at each time point, serially diluted, and spread on AAM agar for viable cell enumeration. The same procedure was repeated with the endospores as inocula. The assay was performed by adding 0.5 mL of the AMPs into 4.5 mL or 4.0 mL AAM broth containing approximately 10^6^ CFU/mL of *A. acidoterrestris* to give final concentrations of 80, 160 μg/mL bovicin HC5, 125, 250 μg/mL of epsilon-polylysine and a combination of both AMPs 80:125, 160:250 μg/mL. Next, the tubes were exposed to thermal shock at 80 °C for 10 min and incubated at 45 °C in a water bath for 0, 12, 24, 36, and 48 h of incubation. The thermal shock at 80 °C for 10 min was repeated after 24, and 48 h of incubation. Aliquots of 100 µL were withdrawn at each time point, serially diluted, and spread on AAM agar for enumeration of viable spores. The number of germinated spores was determined by the difference between the initial and the final spore count at each time point^15^. All tests were performed in technical triplicate and biological duplicates. In every batch, there were four side-by-side treatments: bovicin HC5, epsilon-polylysine, epsilon-polylysine + bovicin HC5, and control (i.e., no antimicrobial peptide added).

### Determination of Decimal reduction time (D)-values for *A. acidoterrestris* DSMZ 2498 endospores in orange juice

The D-value of *A. acidoterrestris* endospores was determined at 95 °C, which is representative of the temperature used during the thermal processing of different fruit juices. Approximately 10^6^ endospores/mL were incubated in glass tubes containing 5 mL of orange juice. The tubes were kept in a water bath at 95 °C and the thermal lag time (approximately 10 min) for the center of the tube to reach the desired temperature was considered. Then bovicin HC5 and epsilon-polylysine at concentrations 80 μg/mL and 125 μg/mL respectively, and also in combination (80:125 μg/mL) were added. For the control (absence of antimicrobial peptides), dilutions (10^-1^ to 10^-6^) of each time sampled were plated, while for the treatments containing the antimicrobial peptides, dilutions (10^-1^ to 10^-6^) at times 0, 2, 4, 6, 8, 10, 12 min were plated. The plating was performed in triplicate using the microdroplet technique, and the plates were incubated at 45 °C for 24 - 48 h. The analysis was carried out in biological duplicates. Survival curves were plotted using the number of germinated endospores surviving in the presence or absence of bovicin HC5 and/or epsilon-polylysine. The data were fitted using linear regression, according to the equation logN = logN_0_ - bt, where N is the number of survivors, N_0_ is the initial number of germinated endospores, t is the exposure time to the treatment temperature, and b is the slope of the survivor curve to reduce one logarithmic cycle. The D-value was calculated by plotting the log of the number of survivors versus time, where D = 1/b, is the inverse of the slope of the survival curve.

### Morphological analysis of the AMPs-treated vegetative cells of *A. acidoterrestris* DSMZ 2498

The effects of combined AMPs on the morphology and cell structure of *A. acidoterrestris* DSMZ 2498 were determined using AFM (Ntegra Prima, NT-MDT Spectrum Instruments; Moscow, Russia). For sample preparation, the cultures of *A. acidoterrestris* DSMZ 2498 were activated in 50 mL of AAM broth, centrifuged (1742 × g for 10 min), and washed with phosphate buffer (pH 4.0, 50 mmol/L) three consecutive times. Subsequently, the pellet was suspended in 20 mL of phosphate buffer (pH 4.0, 50 mmol/L). Then 4.0 or 4.5 mL of the suspension were dispensed in sterilized bottles containing 0.5 mL of each AMP to give final concentrations of 1X and 2X MIC individual AMPs and combined AMPs. Suspension without both AMPs was used as control and incubated for up to 24 h at 45 °C in an electrostatic water bath. Aliquots of 1 mL each were withdrawn after 0, 8, and 24 h of incubation. The samples were centrifuged, the supernatant was discarded, and the cells were spread on glass slides (1 cm × 1 cm) using pipette tips and a drop of sterilized distilled water. These procedures were performed aseptically. The NT-MDT Nova (version 1.26.0.1443) software was used for image acquisition and topography measurements of the *A. acidoterrestris* DSMZ 2498 samples were performed using the intermittent mode to minimize the risk of deformation of the sample and to allow a greater lateral resolution as described by Ribeiro et al.^1^.

### Statistical analysis

All incubations were carried out at least with two technical replicates and the results represent the mean ± standard deviation of two independent experiments. The means were evaluated by one-way ANOVA and Tukey’s post hoc analysis (Graphpad Prism Software, San Diego, California, USA) was used to determine the statistical difference between treatments. A P-value of 0.05 was used to declare significant differences^13^.

## RESULTS

### Effect of AMPs on the growth of *A. acidoterrestris* DSMZ 2498

*A. acidoterrestris* DSMZ 2498, with an initial inoculum of 10^6^ CFU/mL of either vegetative cells or activated endospores, demonstrated an average maximum specific growth rate of approximately 0.26 h^-1^ in AAM (pH 4.0). Inoculation with vegetative cells resulted in a 1-hour lag phase, whereas activated endospores induced a 4-hour lag. With the addition of bovicin HC5, complete growth inhibition was observed at higher concentrations (≥ 2.50 μg/mL) (Figure 1A). The optical densities (OD600 nm) of the culture at all concentrations differ significantly when compared to the control (P < 0.05) after 24 h incubation (Figure 1B). With endospores, growth was completely hindered at ≥ 5.0 μg/mL and an extended lag phase of 14h was observed with concentration 2.50 μg/mL (Figure 2A). The OD600nm of cultures with ≥ 2.50 μg/mL of bovicin HC5 concentrations were significantly different (P < 0.05) from control after 24 h of incubation (Figure 2B).

**Figure 1:**
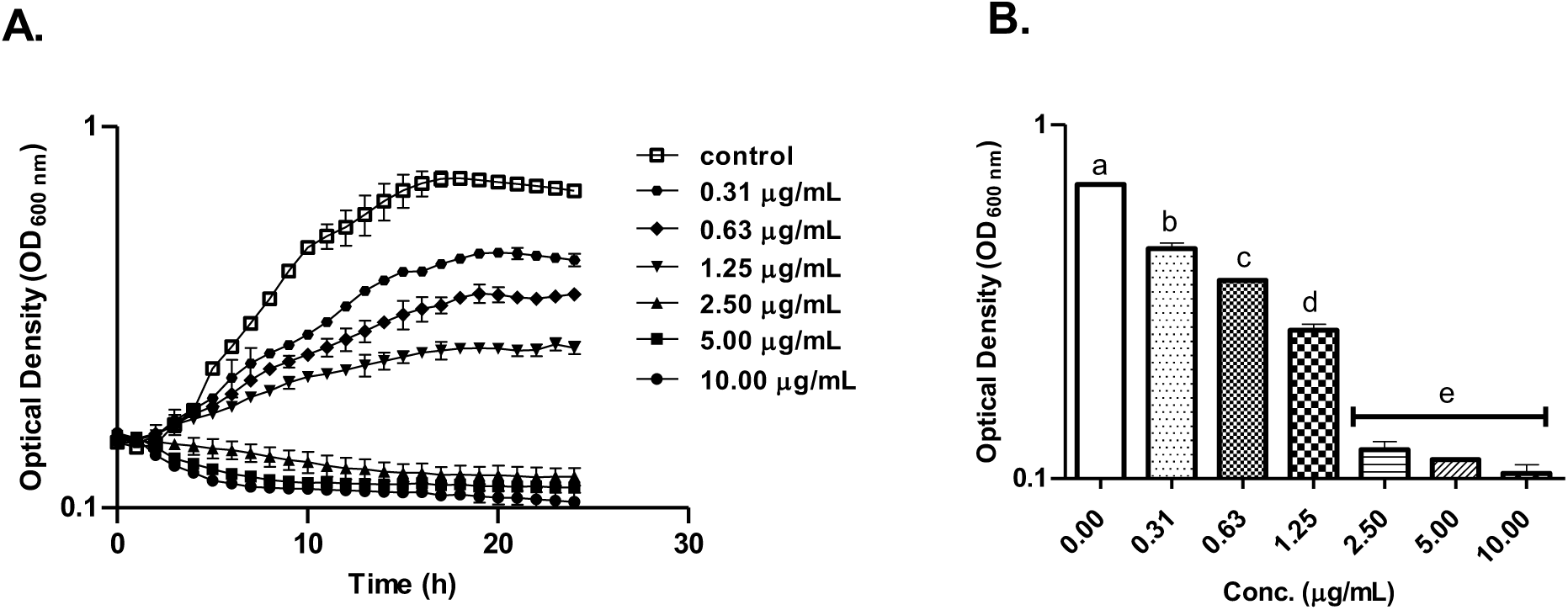
Effect of bovicin HC5 on vegetative cells of *A. acidoterrestris* DSMZ 2498 grown in AAM broth. (A) Growth kinetics of the vegetative cells of *A. acidoterrestris* DSMZ 2498 with different concentrations of bovicin HC5 during 24 h incubation and their (B) optical densities after 24 h incubation. Data were represented as mean ± SD. Bars bearing different letters are significantly different at P < 0.05, while bars bearing the same letter(s) are not significantly different (P > 0.05). Data were analyzed using one-way ANOVA followed by the Tukey post hoc test.

**Figure 2:**
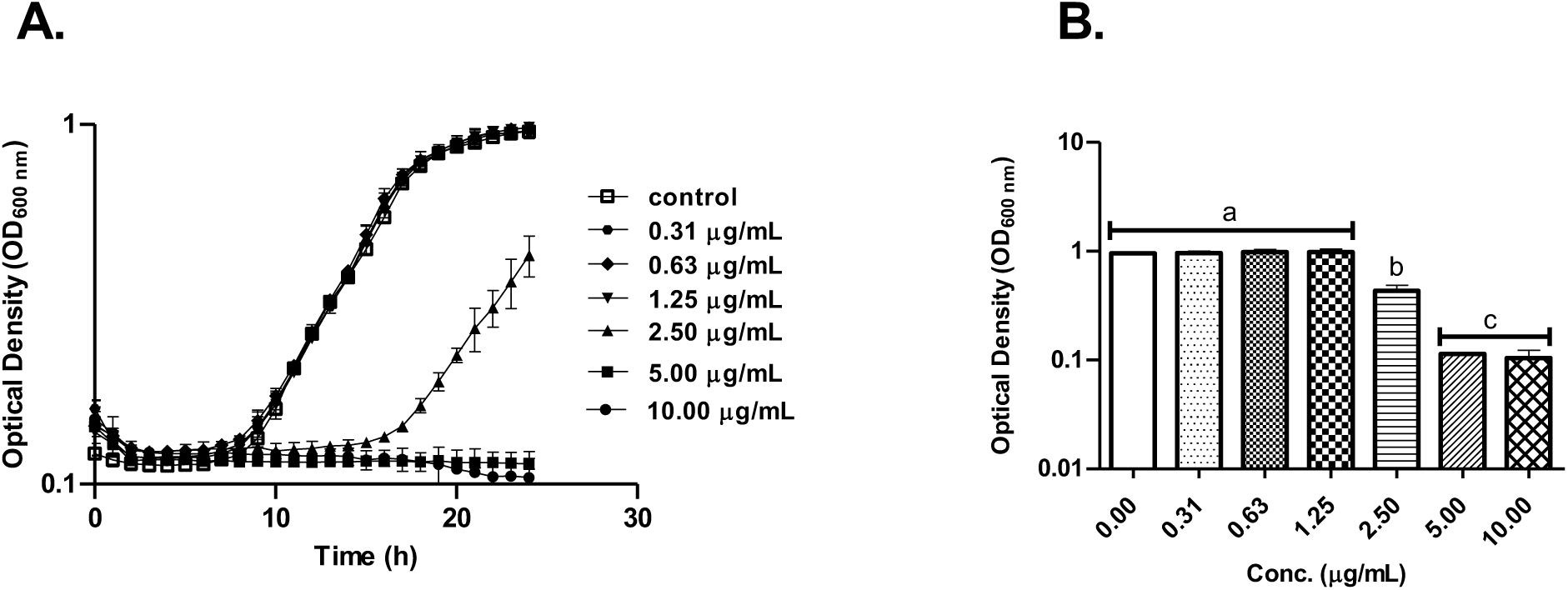
Effect of bovicin HC5 on activated endospores of *A. acidoterrestris* DSMZ 2498 grown in AAM broth. (A) Growth kinetics of germinated endospores of *A. acidoterrestris* DSMZ 2498 with different concentrations of bovicin HC5 during a 24 h incubation and their (B) optical densities after 24 h incubation. Data were represented as mean ± SD. Bars bearing different letters are significantly different at P < 0.05, while bars bearing the same letter(s) are not significantly different (P > 0.05). Data were analyzed using one-way ANOVA followed by the Tukey post hoc test.

With addition of epsilon-polylysine, complete growth inhibition was observed at ≥ 7.81 μg/mL (P < 0.05) with vegetative cells as inocula (Figure 3A). The OD600 nm of all concentrations differed significantly from the control (P <0.05) (Figure 3B). When endospores were used as inocula, no growth was observed with concentrations ≥ 7.81 μg/mL while 3.91μg/mL concentration of epsilon-polylysine exhibited an extended lag phase of 18 h compared to control (Figure 4A). The OD600 nm of the culture at all concentrations of epsilon- polylysine differed significantly (P < 0.05) from the control after 24 h of incubation (Figure 4B).

**Figure 3:**
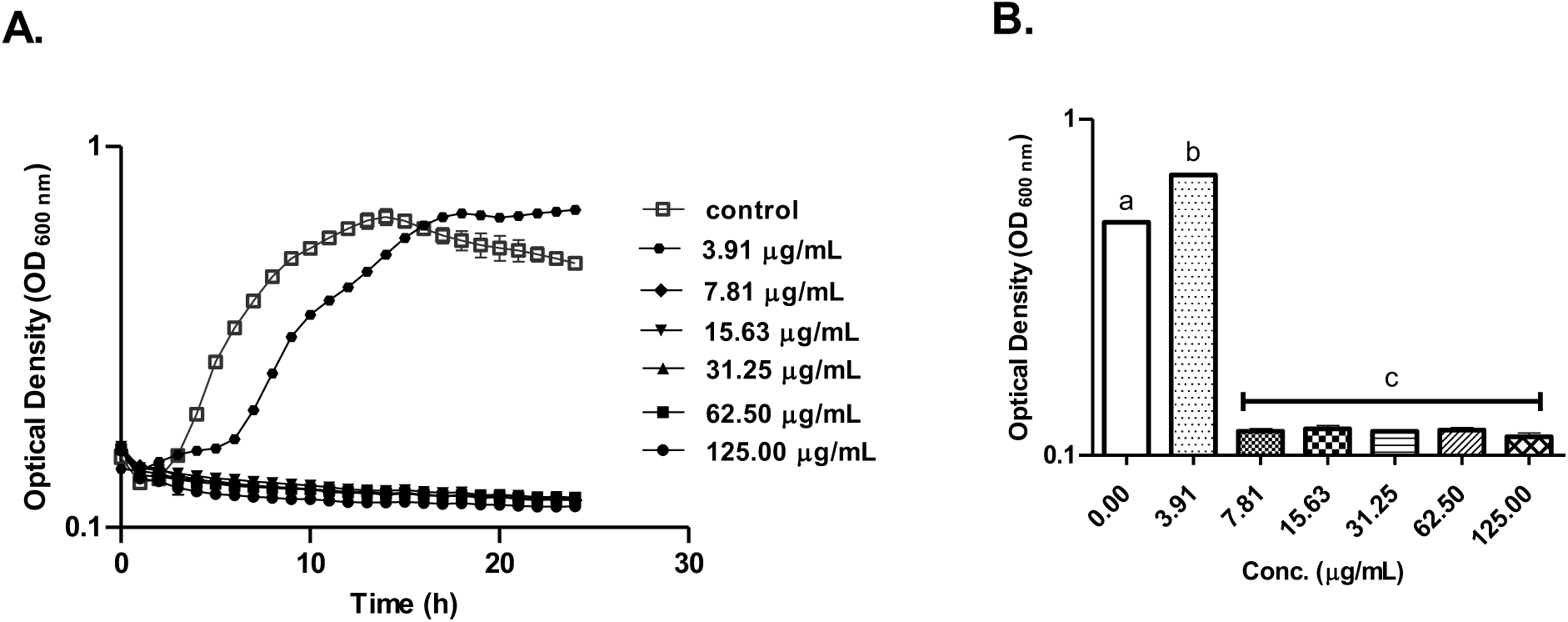
Effect of epsilon-polylysine on vegetative cells of *A. acidoterrestris* DSMZ 2498 grown in AAM broth. (A) Growth kinetics of the vegetative cells of *A. acidoterrestris* DSMZ 2498 with different concentrations of epsilon-polylysine during 24 h incubation and their (B) optical densities after 24 h incubation. Data were represented as mean ± SD. Bars bearing different letters are significantly different at P < 0 .05, while bars bearing the same letter(s) are not significantly different (P > 0.05). Data were analyzed using one-way ANOVA followed by the Tukey post hoc test.

**Figure 4:**
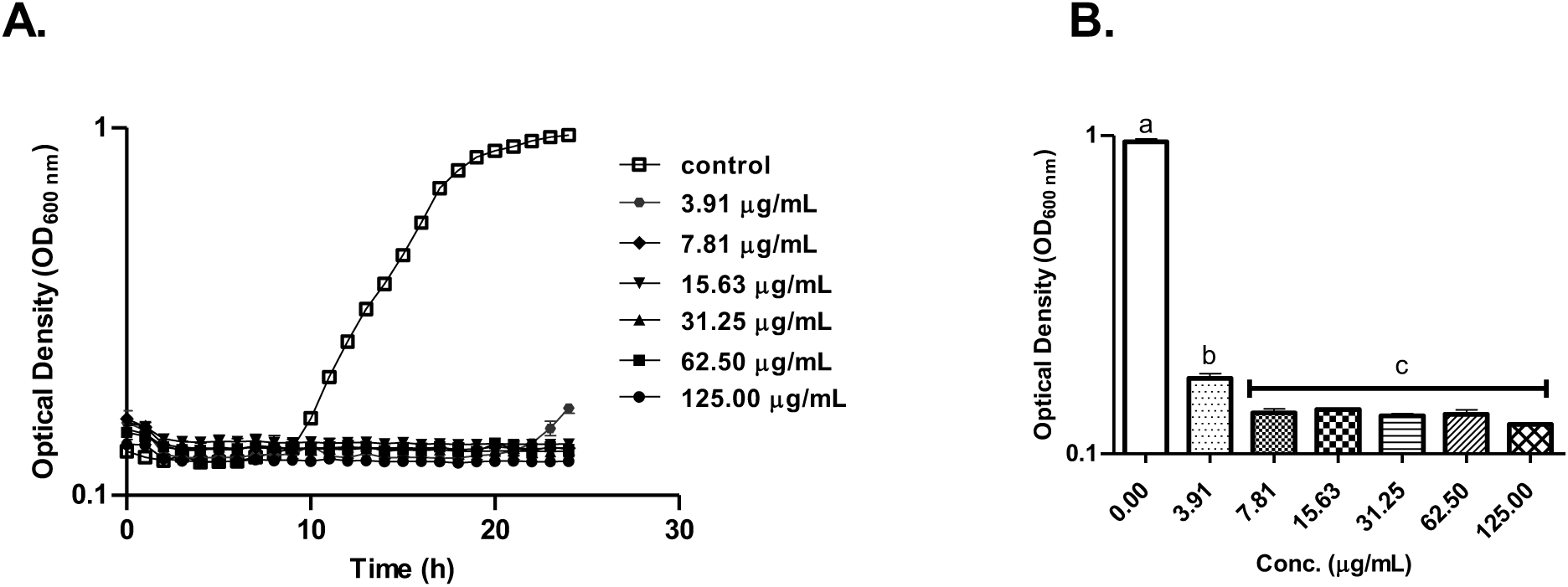
Effect of epsilon-polylysine on activated endospores of *A. acidoterrestris* DSMZ 2498 grown in AAM broth. (A) Growth kinetics of germinated endospores of *A. acidoterrestris* DSMZ 2498 with different concentrations of epsilon-polylysine during a 24 h incubation and their (B) optical densities after 24 h incubation. Data were represented as mean ± SD. Bars bearing different letters are significantly different at P < 0.05, while bars bearing the same letter(s) are not significantly different (P > 0.05). Data were analyzed using one-way ANOVA followed by the Tukey post hoc test.

The MICs for bovicin HC5 were 2.50 μg/mL (vegetative cells) and 5.00 μg/mL (activated endospores), while epsilon-polylysine had a MIC of 7.81 μg/mL for both. The MBC of bovicin HC5 was 5.00 μg/mL for both forms of inocula while epsilon-polylysine showed bacteriostatic effects against vegetative cells and activated endospores across all concentrations (7.81 – 125.00 μg/mL).

### Fractional Inhibitory Concentration (FIC) index of the Combined AMPs

The FIC index of combined AMPs (cAMPs) against *A. acidoterrestris* DSMZ 2498 vegetative cells was based on MIC values for each molecule in combination. The MIC_A_ combined (bovicin HC5 combined with epsilon-polylysine) yielded a MIC of 1.25 μg/mL, compared to MIC_A_ alone (bovicin HC5 alone) at 2.50 μg/mL, resulting in a FIC of 0.5 μg/mL. MIC_B_ combined (epsilon- polylysine combined with bovicin HC5) yielded a MIC of 3.91 μg/mL, compared to MIC_B_ alone (epsilon-polylysine) at 7.81 μg/mL resulting in a FIC of 0.5 μg/mL. The FIC index of the combined AMPs was 1.0, indicating an additive effect.

The FIC index against activated endospores *A. acidoterrestris* DSMZ 2498 followed a similar pattern. MIC_A_ combined resulted in a MIC of 2.50 μg/mL, compared to MIC_A_ alone (5.00 μg/mL), yielding a FIC of 0.5 μg/mL while MIC_B_ combined yielded a MIC of 3.91 μg/mL, compared to MIC_B_ alone of 7.81 μg/mL. yielding a FIC of 0.5 μg/mL The FIC index of the combined AMPs was 1.0, indicating an additive effect.

### Effect of AMPs on the survival of *A. acidoterrestris* in orange juice

In sterilized 100% orange juice, *A. acidoterrestris* DSMZ 2498 (initial inoculum ∼ 6 log CFU/mL) showed consistent viable cell counts after 48 h at 45 °C without AMPs. With addition of bovicin HC5 at 5.0 μg/mL and 10.0 μg/mL, reduced counts were observed at 36 h, followed by a slight increase at 48 h. Higher concentrations of 20.0 - 80.0 μg/mL resulted in varying degrees of reduction, with the latter maintaining a constant decrease at all timepoints (Figure 5A). The area under the time-kill curve (AUC) was significantly reduced for all bovicin HC5 treatments (P < 0.05) (Figure 5B). Addition of epsilon-polylysine at varying concentrations reduced cell counts (4.81 – 4.90 log CFU/mL) compared to that of the control (5.54 log CFU/mL), with 250.0 μg/mL having the most significant effect (Figure 5C). The AUC for epsilon-polylysine treatments was significantly reduced compared to control (P < 0.05) except for 31.25 μg/mL and 62.50 μg/mL (P> 0.05) (Figure 5D). Combined bovicin HC5 and epsilon- polylysine (40.0:62.5 μg/mL and 80.0:125.0 μg/mL, respectively) showed a consistent reduction in viable cell counts at all intervals, reaching below detection limits after 36 h (Figure 5E). AUC for all combinations was significantly different from the control after 48 h (Figure 5F).

**Figure 5:**
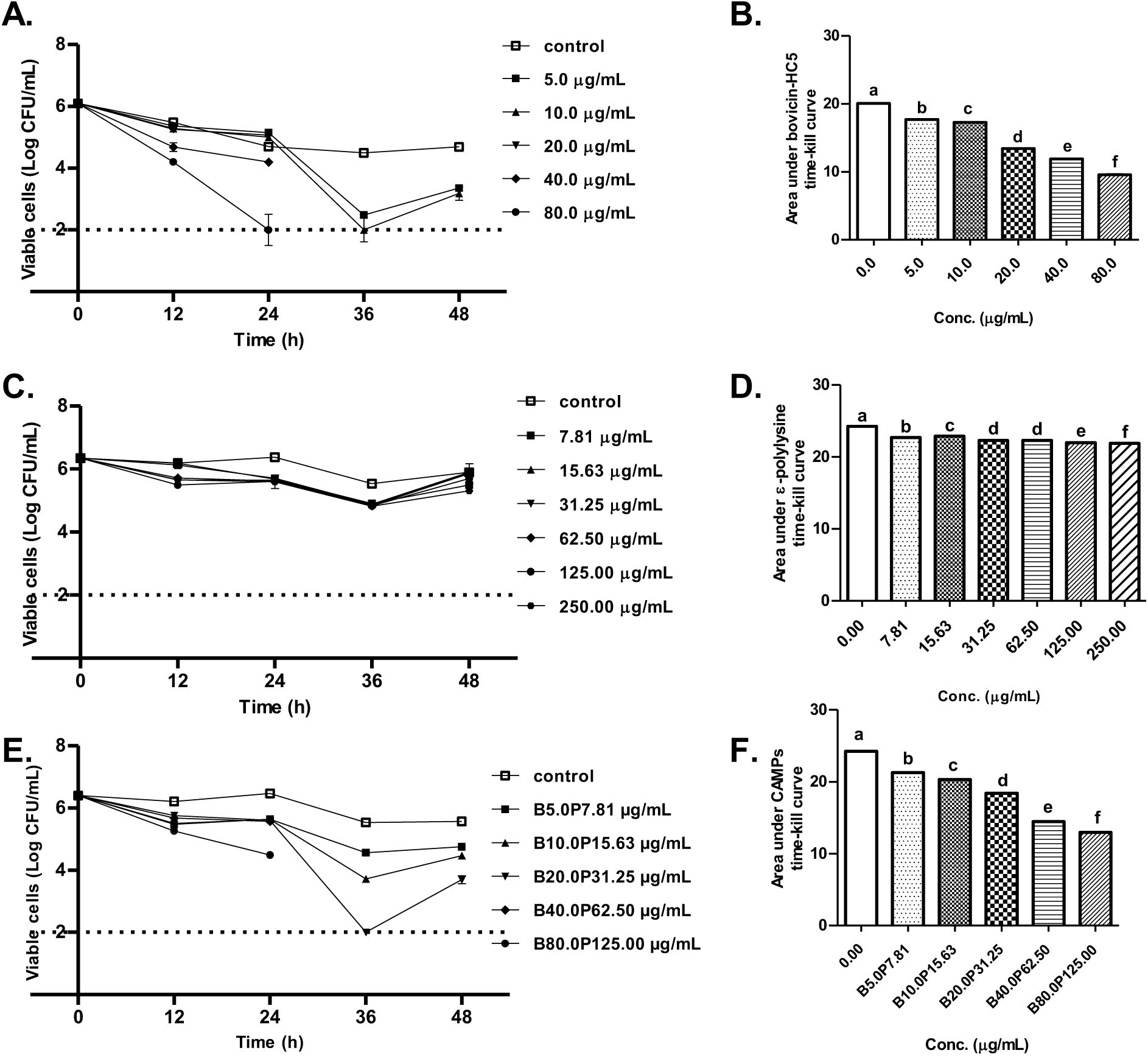
Time-kill curve and AUC of *A. acidoterrestris* DSMZ 2498 (vegetative cells) treated with bovicin HC5 alone (A, B), epsilon-polylysine alone (C, D), and AMPs in combination (E, F) with B, P representing bovicin HC5 and epsilon-polylysine respectively. Data are presented as mean ± SEM, n = 3. Bars showing different letters differ (P < 0.05) from the control. Data were analyzed using one-way ANOVA followed by Tukey post hoc test. Dotted lines represent the detection limit of the assay.

When the juice was inoculated with ∼ 6 log CFU/mL of activated endospores, in the absence of AMPs, viable spore counts decreased by approximately one log cycle. Addition of bovicin HC5 at 80.0 μg/mL and 160.0 μg/mL reduced viable spore counts below detection limits after 36 h (Figure 6A), with significantly reduced AUCs compared to the control (P < 0.05) (Figure 6B). epsilon-polylysine at various concentrations reduced viable spore counts (3.84 - 3.60 log CFU/mL) compared to that of control, 4.30 log CFU/mL) (Figure 6C), with significant differences in AUCs (P < 0.05) (Figure 6D). Combined AMPs consistently reduced viable spore counts below detection limits after 36 h (Figure 6E), and also significantly reduced AUCs compared to the control (P < 0.05) (Figure 6F).

**Figure 6:**
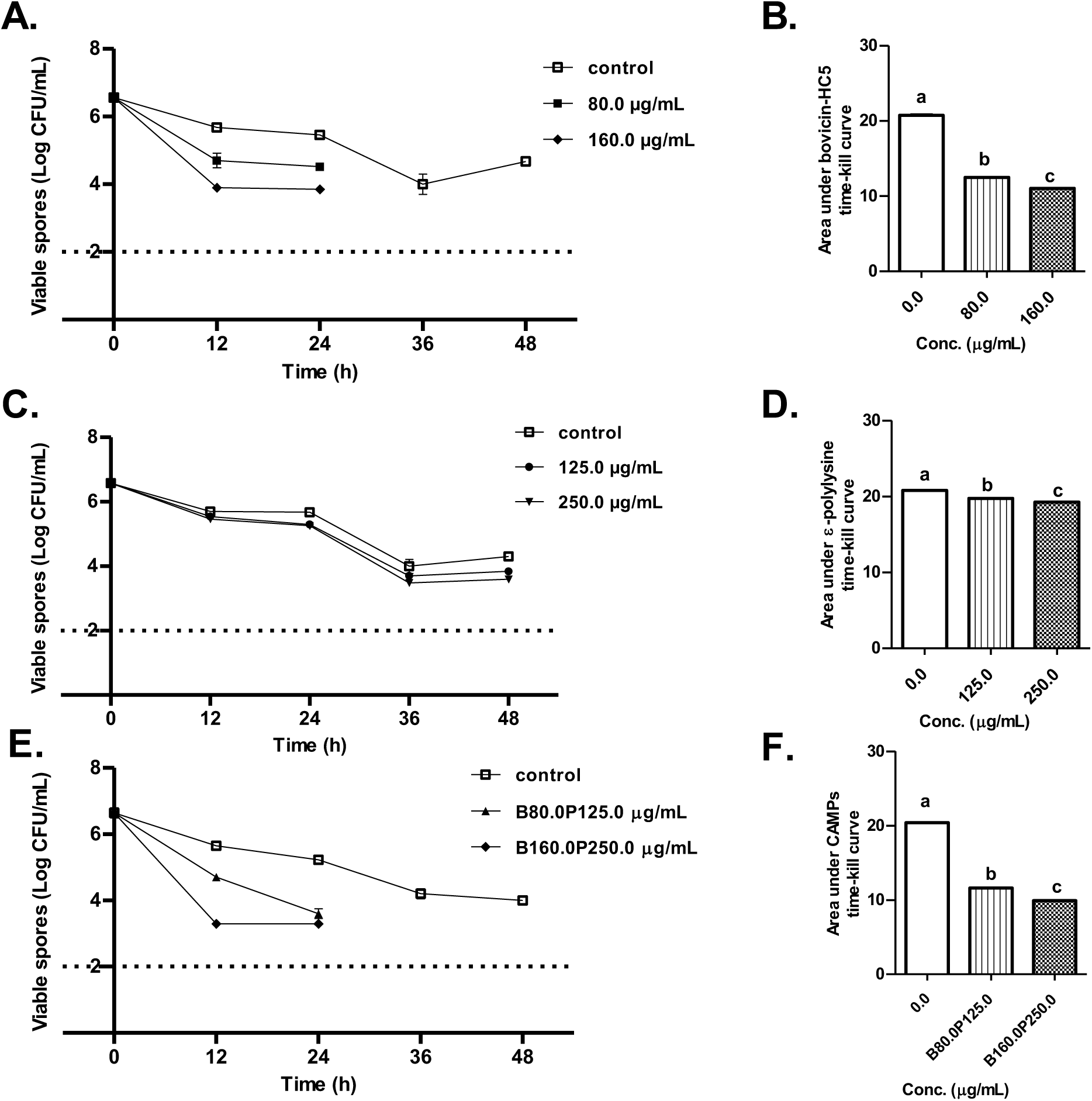
Time-kill curve and AUC of *A. acidoterrestris* DSMZ 2498 germinated endospores in orange juice and treated with bovicin HC5 alone (A,B), epsilon-polylysine alone (C,D), and AMPs in combination (E,F) with B, P representing bovicin HC5 and epsilon-polylysine respectively. . Data are presented as mean ± SEM, n = 3. Bars showing different letters are significantly different (P < 0.05) from control. Data were analyzed using one-way ANOVA followed by Tukey post hoc test. Dotted lines represent the detection limit of the assay.

### Effect of the AMPs on the thermal resistance of *A. acidoterrestris* endospores in orange juice

The average D-value at 95 °C (D95 °C) of *A. acidoterrestris* DSMZ 2498 endospores in orange juice without AMPs was 7.68 min. The addition of bovicin HC5 (80 µg/mL) decreased (P < 0.05) the D95 °C value to 2.19 min, which represented a 71.48 % reduction in the number of viable spores compared to the control (Figure 7A, Table 1). In contrast, 125 µg/mL of epsilon- polylysine in orange juice with endospores showed no significant difference (P > 0.05) from the control after 12 min of heating (Figure 7B, Table 1). Orange juice containing bovicin HC5 and epsilon-polylysine at 80:125 µg/mL, respectively showed a significant reduction in the number of viable spores below the detection limit (P < 0.05) with a D95 °C of 2.43 min. Therefore, addition of cAMPs to orange juice decreased the D95 °C of *A. acidoterrestris* DSMZ 2498 endospores by 68.36 %, close to the effect observed when bovicin HC5 alone was added to orange juice (Figure 7C, Table1).

**Figure 7:**
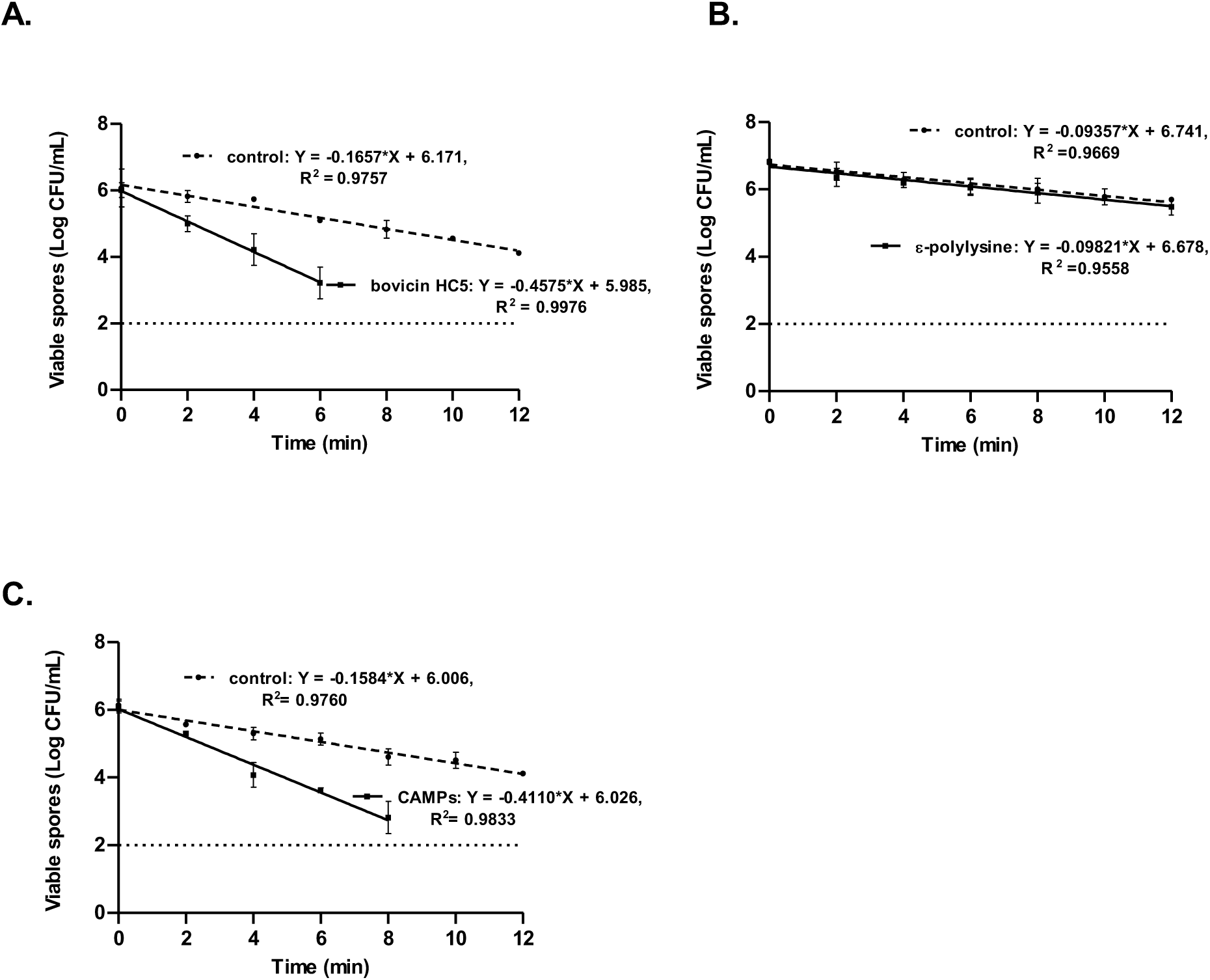
Survival curve of *A. acidoterrestris* DSMZ 2498 endospores in orange juice. The orange juice was treated with bovicin HC5 at 80 µg/mL (A), epsilon-polylysine at 125 µg/mL (B), or a combination of both AMPs (B80:P125 μg/mL) (C). The incubation was carried out at a temperature of 95 °C for 12 min. The dotted line represents the detection limit.

**Table 1:** Effect of bovicin HC5, epsilon-polylysine or both AMPs in the D-value at 95 °C of *A. acidoterrestris* DMSZ 2498 endospores in orange juice.

| AMPs | Concentration (µg/mL) | D-value (min)* | Reduction (%) |
| --- | --- | --- | --- |
| Bovicin HC5 | 80.0 | 2.19±0.35 | 71.48 |
| Epsilon-Polylysine | 125.0 | 7.31±1.41 | 4.78 |
| cAMPs | 80.0:125.0 | 2.43±0.23 | 68.36 |

D-value of control = $7.68 \pm 1.51$ min. \*Values are mean ± standard error of mean (n=3)

### Morphological characteristics of *A. acidoterrestris* DSMZ 2498 treated with AMPs

Untreated vegetative cells of *A. acidoterrestris* DSMZ 2498 exhibited regular, smooth, and fully rounded rod shape even after 24 h of incubation in phosphate buffer (pH 4) at 45 °C. However, upon treatment with the antimicrobial peptides, distinct changes in the cell morphology were observed through atomic force microscopy (AFM) (Figures 8, 9, 10).

**Figure 8:**
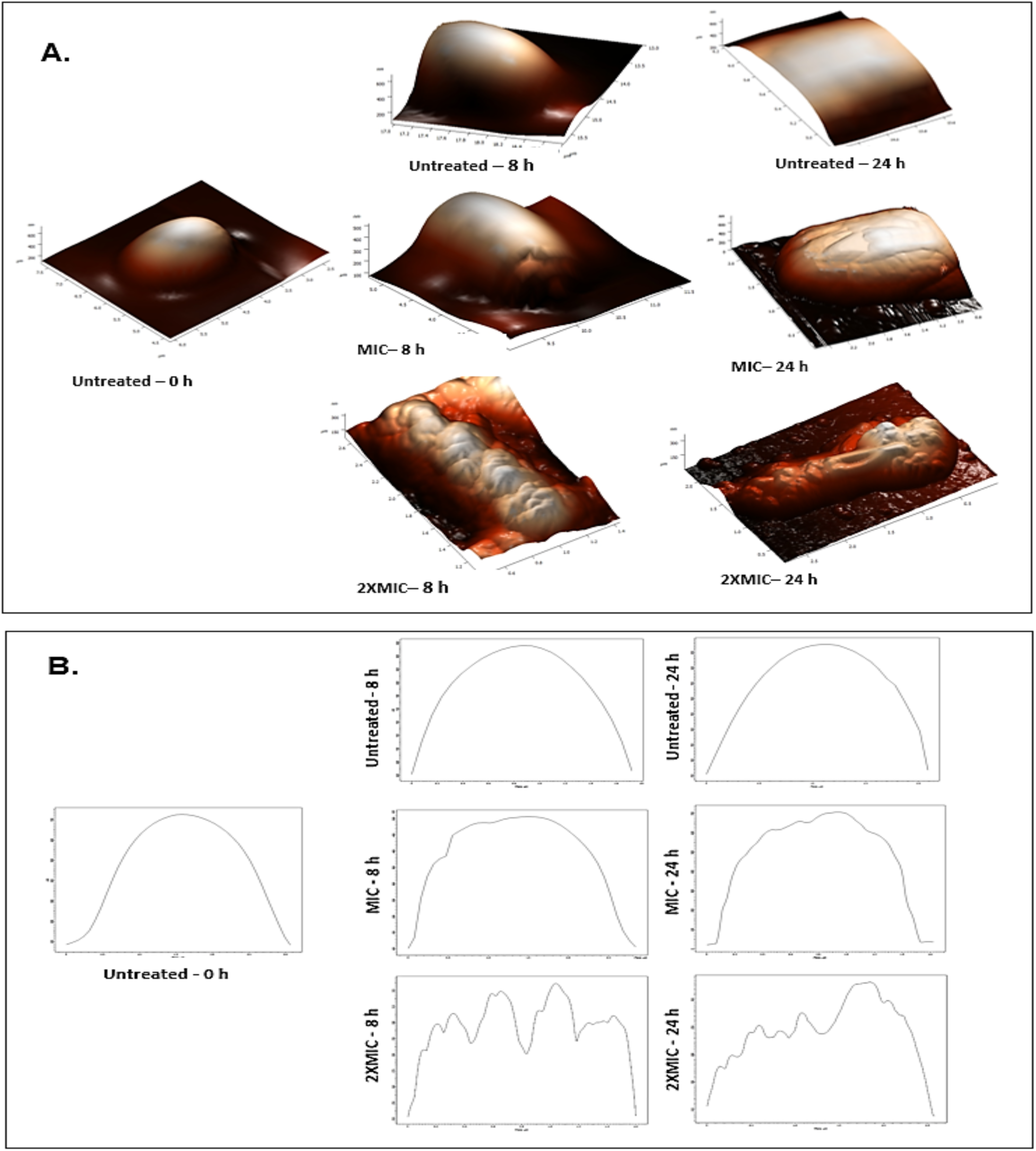
Effect of bovicin HC5 on vegetative cells of *A. acidoterrestris* DMSZ 2498. Three-dimensional topographical images (Panel A) and cross-sectional measurements (Panel B) of vegetative cells of *A. acidoterrestris* DMSZ 2498 obtained by AFM (NT-MDT) after 0, 8 h, 24 h of incubation. The cell suspensions were incubated with MIC - 2.5 µg/mL and 2XMIC - 5.0 µg/mL of bovicin HC5 in phosphate buffer (pH 4.0) at 45 °C. The images were observed using a scanning area of 1 to 4 μm.

Treatment with 2.5 and 5.0 μg/mL (1X and 2X MIC) of bovicin HC5 resulted in rough areas and deformation of the cell surface with incubation time. Cultures treated with 5.0 μg/mL of bovicin HC5 exhibited greater deformation, with wrinkles observed after 8 h of incubation and the presence of deep dents in the center of cells after 24 h (Figure 8). When 7.81 and 15.63 μg/mL (1X and 2X MIC) of epsilon-polylysine were used against the vegetative cells, irregular cell surface and shape deformation were observed after all incubation times (Figure 9).

**Figure 9:**
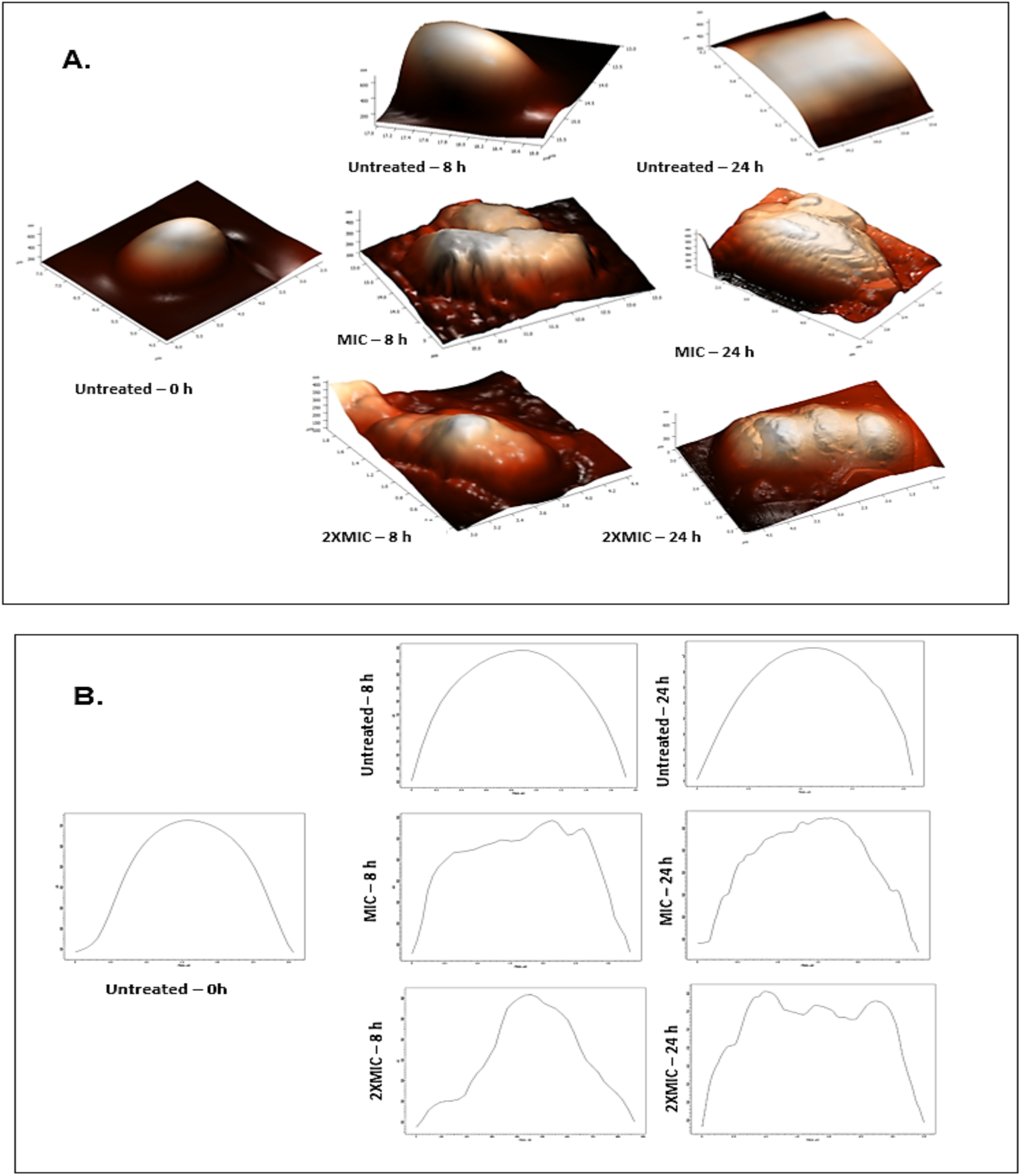
Effect of epsilon-polylysine on vegetative cells of *A. acidoterrestris* DMSZ 2498. Three-dimensional topographical images (Panel A) and cross-sectional measurements (Panel B) of vegetative cells of *A. acidoterrestris* DMSZ 2498 obtained by AFM (NT-MDT) after 0, 8 h, 24 h of incubation. The cell suspensions were incubated with MIC - 7.81 µg/mL and 2XMIC -15.63 µg/mL of bovicin HC5 in phosphate buffer (pH 4.0) at 45 °C. The images were observed using a scanning area of 1 to 4 μm.

Furthermore, treatment with 1X and 2X MIC of combined bovicin HC5 and epsilon-polylysine (1.25:3.91 μg/mL and 2.50:7.81 μg/mL, respectively) led to increased cell surface roughness and the presence of extrusions after each incubation time point (Figure 10).

**Figure 10:**
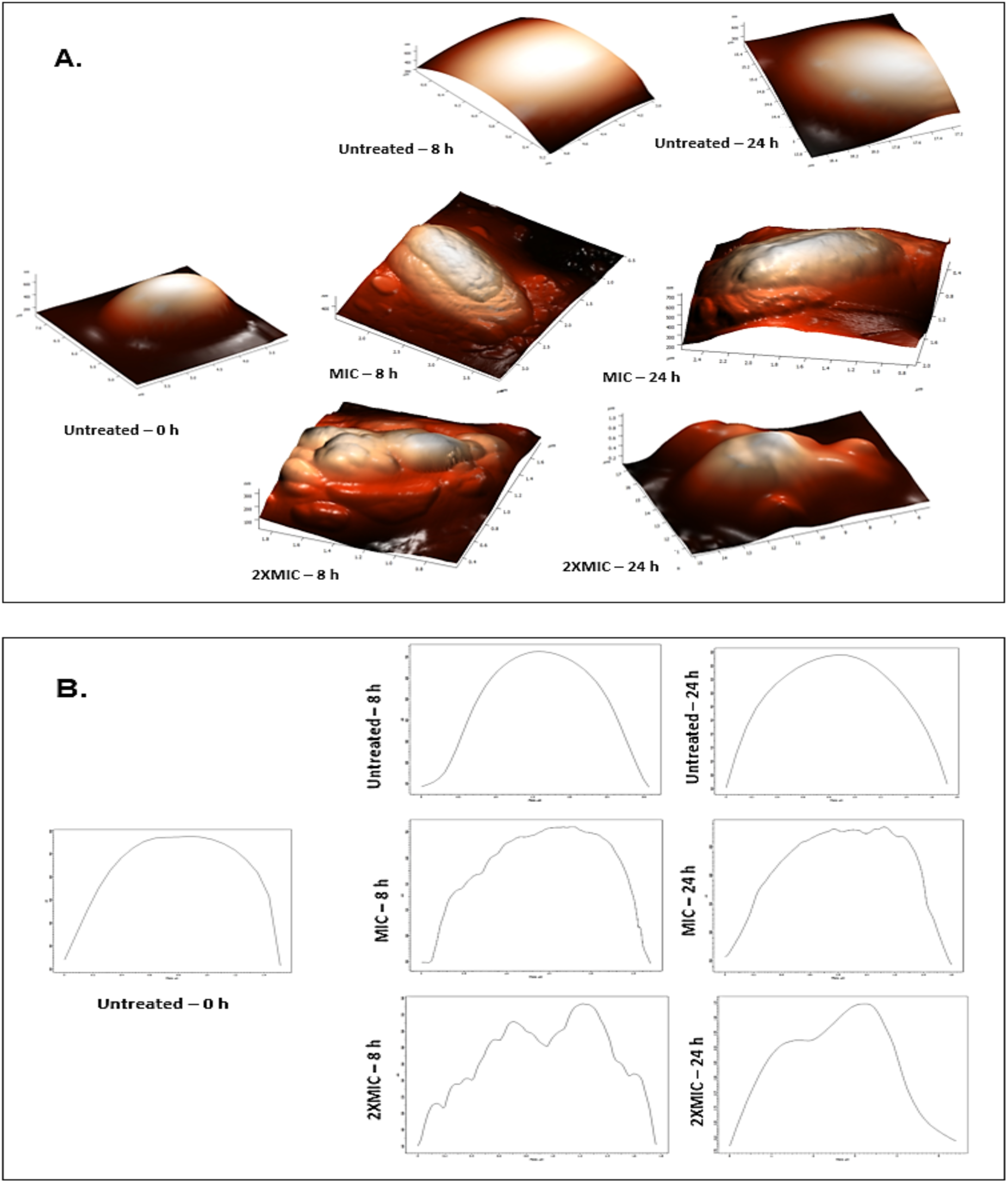
Effect of combination of bovicin HC5 and epsilon-polylysine on vegetative cells of *A. acidoterrestris* DMSZ 2498. Three-dimensional topographical images (Panel A) and cross-sectional measurements (Panel B) of vegetative cells of *A. acidoterrestris* DMSZ 2498 obtained by AFM (NT-MDT) after 0, 8 h, 24 h of incubation. The cell suspensions were incubated with MIC - 1.25:3.91 µg/mL and 2XMIC - 2.50:7.81 µg/mL of the combination of bovicin HC5 and epsilon-polylysine in phosphate buffer (pH 4.0) at 45 °C. The images were observed using a scanning area of 1 to 4 μm.

## DISCUSSION

*A. acidoterrestris* has been responsible for several spoilage incidents associated with unpleasant changes in the odor and taste of fruit juices due to the production of spoilage compounds such as guaiacol^7^. Given that traditional food processing methods have limited efficacy to prevent this spoilage of acidic beverages by *A. acidoterrestris,* it is of most importance to study alternative methods to improve the safety of these products^14^.

In this study, the effect of antimicrobial peptides on the growth of vegetative cells and germination of endospores *A. acidoterrestris* were investigated in AAM broth. Based on the results obtained, bovicin HC5 and epsilon-polylysine showed effective antimicrobial effects and caused varying reductions in growth rate and lag phase of *A. acidoterrestris*. The noticeable variation in the growth inhibition of *A. acidoterretris* when different inocula (vegetative cells and endospores) were exposed to 2.50 μg/mL of bovicin HC5 suggests that the bovicin HC5 may interfere with spore germination, though not sufficient to completely inhibit growth in comparison to that of the vegetative cells at the same concentration. According to Gut et al.^22^, one possible explanation is that, like nisin, binding of bovicin HC5 to lipid II (achieved with the above-mentioned concentration) can inhibit the vegetative cells but is inadequate to effectively inhibit spore outgrowth. Similar to other lantibiotics, bovicin HC5 exhibits a higher affinity for lipid II than anionic lipid membranes^23^. This may help to explain why the growth rate of *A. acidoterrestris* germinated endospores with bovicin HC5 concentrations ≤ 2.50 μg/mL was higher than that of vegetative cells. More bovicin HC5 has likely been bound to lipid II on the spore membrane, rendering it unavailable to bind to the vegetative cells that emerged from the germinated spores. However, further studies will be required to fully understand the underlying mechanism. The strong poly-cationic property of epsilon-polylysine, which supports its capacity to adhere to anionic surfaces, thus increasing cell membrane permeability and causing inhibition of enzyme activity, may be responsible for its inhibitory action^10^. When both inocula (vegetative cells and endospores) of *A. acidoterrestris* were used in the presence of 3.91 μg/mL epsilon- polylysine, variations in the growth rate and lag phase of *A. acidoterrestris* were observed. This may be due to the fact that during spore germination the spore membrane becomes permeable^24^, which further aids the membrane permeability of epsilon-polylysine and facilitates its penetration into the germinated spore.

Current research demonstrated several benefits of antimicrobial peptides over chemical preservatives, suggesting that these molecules could enhance the storage life of food products by preventing the growth of food-spoilage microorganisms. Hence, by combining two antimicrobial peptides, the efficacy of antimicrobial peptides could be increased, resulting in additional hurdle against food spoilage^25^. In this study, the combined effect of bovicin HC5 and epsilon-polylysine against vegetative cells and germinated endospores of *A. acidoterrestris* DSMZ 2498 proved to be additive (FIC index = 1) in AAM broth at pH 4, which implies that the combined effect is the sum of the effect of the two antimicrobial peptides working independently. Liu et al.^26^ reported that although bacteriocins alone have effective antimicrobial properties, their use in combination with other antimicrobials hurdles promotes additive/synergistic effects. For a synergistic effect, it is suggested that mechanistic insights associated with the combined effects of the antimicrobials in different substrates will be helpful to achieve optimal results^26^.

Furthermore, the time-kill assays demonstrated significant inhibitory effects on the growth of vegetative cells and spore germination of *A. acidoterrestris* after exposure to both AMPs (individually and in combination) in orange juice. Both AMPs replicated their individual bacteriostatic or bactericidal activities in orange juice as observed in the laboratory media but at higher concentrations. These results suggest the interaction of this bacteriocin with components of the fruit juice matrix^27^. However, bovicin HC5 showed a more pronounced antibacterial effect compared to that of epsilon-polylysine.

To implement effective strategies for fruit juice preservation, evaluation of the heat resistance in spoilage-causing microorganisms is important^1^. The heat resistance of *A. acidoterrestris* inoculated in orange juice obtained in this study was represented by a mean D-value of 7.68 min at 95 °C. It is important to note that the D-value of the endospores of *A. acidoterrestris* is affected by varying factors such as growth conditions or characteristics of the study design, including food matrix used (i.e, type of fruit juice), media or fruit juice pH, TSS content of fruit juices, temperature, quality of the culture medium, inactivation method, bacterial strain tested, among others^28^. For different combinations of these factors, Huertas et al.^29^ reported a D95 °C value of 5.24 min of *A. acidoterrestris* in orange juice. Comparably, these results confirm the high heat resistance of *A. acidoterrestris* endospores and the ineffectiveness of pasteurization with a temperature range of 80 – 100 °C for less than 30 s, commonly used in fruit juices processing^30^.

The efficacy of bovicin HC5 (individually and in combination) to significantly reduce the heat resistance of endospores of *A. acidoterrestris* in this study can be attributed to its high stability under pasteurization temperatures^1^. Although its sporicidal effect has not been completely elucidated, its interaction with endospores could be similar to what has been reported for nisin^24^. The effect of bovicin HC5 on endospores might occur after germination has been initiated. When endospores germinate, lantibiotics such bovicin HC5 could bind to the lipid II in the cell membrane thus inhibiting the outgrowth of endospores and reducing viability^24^. The reduction in effect of epsilon-polylysine when added to orange juice at pH 3.68 at a temperature of 95 °C could be attributed to its amine groups which are strongly cationic at pH values < 9.0 allowing it to interact with anionic components within the food matrix, forming insoluble precipitates which further affects its ability to interact with the bacterial cell membrane^31^. Also, this result reveals the need to investigate other varying factors that affect the activity of epsilon-polylysine under food processing conditions.

Normal cell morphology and an integral cell membrane are vital for bacterial growth and maintenance of vital metabolic functions^32^. Previous studies demonstrated that some antimicrobial peptides like enterocin AS-48, bificin C6165, bacteriocin RC20975, nisin, and bacteriocin AMA-K could induce structural changes in *A. acidoterrestris* cells^33–36^. In the current study, both epsilon-polylysine and bovicin HC5 as well as their combinations at MIC concentration and 2XMIC induced drastic changes in the cell morphology of *A. acidoterrestris*. The structural changes observed on the vegetative cells of *A. acidoterrestris* treated with bovicin HC5 provides further evidence of its dual mechanism of action through inhibition of cell wall biosynthesis and formation of pores in the cell membrane, resulting in cell death as also reported by Ribeiro et al.^1^. Additionally, the pore-like lesions observed provide evidence of depolarization of the membrane potential and cellular leakage, which eventually renders the bacterial cell non-viable as described by Egan et al.^24^. The effects of epsilon-polylysine are similar to that reported by Tan et al.^10^ against *S. aureus*. The authors demonstrated that epsilon-polylysine induced structural change of the peptidoglycan in the cell wall and affected the integrity of the cytoplasmic membrane in treated cells. The effect of the combined MICs of epsilon-polylysine and bovicin HC5 which were a combination of 50 % lower concentrations of the MICs of the individual AMPs revealed structural changes similar to those obtained with the MICs of the AMPs and this result was replicated with 2XMIC as shown in Figure 14. These results correlate with the Loewe additivity theory, the effect of antimicrobials in combination is determined not by the sum of their normalized effects, but rather by the sum of their normalized dosages, such that their combined effect is the same across all combinations that have the same total normalized dosage^37^.

Overall, the findings in this study indicate that bovicin HC5 and epsilon-polylysinecould be used as natural preservatives in hurdle technologies to control *A. acidoterrestris* and improve the microbiological stability and safety of fruit juices.

## ACKNOWLEDGEMENTS

The author thanks the Coordenação de Aperfeiçoamento de Pes-soal de Nível Superior (CAPES), Brasília, Brazil, and the Tertiary Education Trust Fund (TETFund), Nigeria in collaboration with the Forum for Agricultural Research in Africa (FARA) for the granting of scholarships and financial support to conduct this research.

